# How large and diverse are field populations of fungal plant pathogens? The case of *Zymoseptoria tritici*

**DOI:** 10.1101/2022.03.13.484150

**Authors:** Bruce A. McDonald, Frederic Suffert, Alessio Bernasconi, Alexey Mikaberidze

## Abstract

Pathogen populations differ in the amount of genetic diversity they contain. Populations carrying higher genetic diversity are thought to have a greater evolutionary potential than populations carrying less diversity. We used published studies to estimate the range of values associated with two critical components of genetic diversity, the number of unique pathogen genotypes and the number of spores produced during an epidemic, for the septoria tritici blotch pathogen *Zymoseptoria tritici*. We found that wheat fields experiencing typical levels of infection are likely to carry between 3.2 and 14.9 million pathogen genotypes per hectare and produce at least 2.3 to 10.5 trillion pycnidiospores per hectare. Given the experimentally derived mutation rate of 3 × 10^−10^ substitutions per site per cell division, we estimate that between 28 and 130 million pathogen spores carrying adaptive mutations to counteract fungicides and resistant cultivars will be produced per hectare during a growing season. This suggests that most of the adaptive mutations that have been observed in *Z. tritici* populations can emerge through local selection from standing genetic variation that already exists within each field. The consequences of these findings for disease management strategies are discussed.

## 1. Introduction

The evolutionary potential of pathogen populations is affected by many factors, including the amount of genetic diversity present in a population and the number of offspring produced during an epidemic. Pathogen population size is known to impact the rates at which genetic diversity is eroded by genetic drift or increased by accumulation of mutations (Charlesworth 2009; Zwart et al. 2011) and is influenced by the reproductive capacity of the pathogen as quantified by the basic reproductive number *R*_0_. While several empirical studies have documented that natural epidemics are caused by plant pathogen populations with different degrees of genotypic diversity, very few sought to estimate the actual number of pathogen genotypes colonizing a crop during a typical epidemic. Many studies have quantified the number of offspring, usually spores, produced *in vitro* by a relatively small number of strains for various species of plant pathogens, but fewer have quantified the number of spores produced by these species *in planta* (mostly in growth chambers or greenhouses under highly standardized environmental conditions). Even fewer studies have estimated the number of pathogen spores produced during natural field epidemics under naturally fluctuating environmental conditions, while the most relevant scale to infer the consequences for the population dynamics and evolutionary potential of crop pathogens is a crop canopy or an entire field.

Spore production has been quantified for many plant pathogens, sometimes by combining experimental and modelling approaches (e.g. Sache and de Vallavieille-Pope 1995; Leclerc et al. 2017). In some pathosystems, estimates of *R*_0_ are of the order of tens (<35 in *Puccinia striiformis* f.sp. *tritici*; Mikaberidze et al. 2016) or even hundreds (>300 in *Puccinia lagenophorae*; Frantzen and van den Bosch et al. 2000), highlighting the potential for rapid multiplication to form massive populations, whereas more moderate estimates are found for the *Zymoseptoria tritici*--wheat pathosystem (<10; Mikaberidze et al. 2017). Sporulation capacity, often estimated by the number of spores produced by a fruiting body at a single point in time, is considered to be a crucial life history trait of a pathogen (e.g., Pariaud et al. 2009; Delmas et al. 2016; Suffert et al. 2013). More accurate assessments of this component take into account the depletion dynamics of the sporulating structures, such as the pustules of rusts or the pycnidia of Ascomycetes, over several points in time (e.g., Sache 1997; Eyal 1971).

The heterothallic ascomycete fungus *Zymoseptoria tritici* causing septoria tritici blotch (STB) on wheat is a particularly relevant pathogen to answer the question posed in our title. The development of STB epidemics is driven by a mixed reproductive system, characterized by both splash-dispersed asexual pycnidiospores released from pycnidia during the wheat growing season and wind-dispersed ascospores released from pseudothecia, the sexual fruiting bodies that are produced on senescent tissues and wheat residues (Suffert et al. 2011; Singh et al. 2021). Lesions can be found at high numbers on living leaf tissues in wheat fields over the majority of the growing season. Asexual fruiting bodies (pycnidia containing pycnidiospores) are numerous within lesions and the capacity of sporulation for each fruiting body is high. The intensity of an epidemic is determined mainly by the number of secondary infections and thus by the total number of pycnidiospores produced. The genetic diversity of a local pathogen population is high due to recurring cycles of sexual reproduction, that occurs mainly between the end of an epidemic season and the beginning of the subsequent growing season, but also during the growing season (Zhan et al. 1998). The recent discovery of chlamydospores in *Z. tritici* (Sardinha-Francisco et al. 2019) adds a new element to the pathogen life cycle that can also contribute to the effective population size and maintenance of genetic diversity by enabling the buildup of a chlamydospore bank in soils planted to wheat and long-term persistence of the pathogen between crop rotations.

The population genetics and population genomics of *Z. tritici* has been well characterized by using transect sampling to obtain well annotated collections of strains coming from naturally infected wheat fields distributed around the world (McDonald and Mundt 2016; Boixel et al. 2022), enabling comparisons of population structure across different geographical scales. In some cases, the same field site was sampled over several years (Zhan et al. 2001; El Chartouni et al. 2012; Morais et al. 2019; Oggenfuss et al. 2021), enabling comparisons of population structure over time. The field populations were characterized using both neutral genetic markers such as RFLPs and SSRs (McDonald and Martinez 1990; Linde et al. 2002; Drabesova et al. 2013) and DNA sequences of strongly selected genes encoding fungicide resistance and avirulence effectors (Boukef et al. 2012; Brunner and McDonald 2018; Brunner et al. 2008). The main outcome of these analyses was to show that field populations around the world typically contain high levels of both gene and genotype diversity. The gene diversity reflects the high effective population size and the genotype diversity reflects recurring cycles of sexual recombination. Comparisons of neutral allele frequencies over different spatial scales indicated that gene flow is high across regional scales of hundreds of km and that populations from different continents are very similar to each other, consistent with significant gene flow occurring among continents, with the notable exception of Australia (Juergens et al. 2006). Measurements of changes in neutral allele frequencies over time combined with field experiments using mark-release-recapture experimental designs provided rough estimates of effective population size and genotype diversity at the field scale (Zhan et al. 2001; Zhan and McDonald 2004). But no attempt was made until now to methodically calculate the amount of genotype diversity and spore production occurring in naturally infected fields.

Here we focus mainly on the asexual stage of STB epidemics. We bring together a wide variety of results from the literature to generate estimates of the number of genotypes found in a hectare of wheat experiencing a natural STB epidemic and also estimate the number of pycnidiospores produced per hectare in fields with different levels of infection. These results combine concepts and results coming from two disciplinary fields: ‘Phytopathometry’ (Bock et al. 2021) and ‘Population genetics’ (Burdon and Laine 2019) that are not often brought together in plant pathology. Based on our analyses, we estimate that wheat fields showing typical levels of STB infection carry at least 3.2 million *Z. tritici* genotypes per hectare and produce at least 2.3 trillion pycnidiospores that can result in 28 million adaptive mutations per hectare. These numbers underline the tremendous evolutionary potential of local pathogen populations and the challenges faced by breeders aiming to create durably resistant wheat cultivars and pesticide manufacturers aiming to avoid the emergence of fungicide resistance.

## 2. Methods

In order to estimate the number of pathogen genotypes and the number of pathogen spores existing in a field, we first need to know the number of infected host units existing in a well-defined spatial context. For STB, the most relevant host unit is an infected leaf and we chose one hectare as the most appropriate spatial scale. To estimate the number of wheat leaves found in a square meter in a typical wheat field, we used information provided in Lloveras et al. (2014). According to that source, northern central European wheat farmers typically aim for a planting density of approx. 200 plants per square meter that provide between 475-500 fertile tillers producing ears (spikes) of grain per square meter. The recommendation in North America is also to aim for 200 plants per square meter, with an increase of up to 300 plants per square meter if the fields are irrigated, with a goal of achieving between 500-600 fertile tillers per square meter. In the drier Mediterranean climate, Lloveras et al. (2014) showed that 400-500 plants were an optimum planting density that would provide between 380-500 fertile tillers per square meter. Based on these reported average values, our calculations assume that a typical wheat field will contain 450 fertile tillers per square meter. If we assume that only the top 4 leaves on each tiller can be infected during the most active phase of an STB epidemic (we recognize that this likely underestimates the actual number of leaves), this would provide an average leaf density of 1800 per square meter, or 18 million leaves per hectare available to be infected by STB. We will use this estimate in our general calculations of genotypic diversity and spore production. Other estimates could be prepared for specific cases where there are lower or higher planting densities according to the local agricultural practices.

To estimate the number of unique *Z. tritici* genotypes found in a field, we used published results coming from naturally infected wheat fields at 31 field sites in nine countries on four continents collected over more than 30 years (Table 1) (McDonald and Martinez 1990; Linde et al. 2002; Juergens et al. 2006; Boukef et al. 2012; Drabešová et al. 2013; Morais et al. 2019). Pathogen genotypes were identified using neutral genetic markers, including restriction fragment length polymorphisms (RFLPs) and microsatellites (SSRs). In these studies, diseased leaves were sampled from the four upper leaf layers and in total, more than 2400 pathogen isolates were genotyped (Table 1). An average of one *Z. tritici* genotype was found per leaf across the field populations, but this estimate is strongly affected by the sampling procedure. In most field collections, only a single isolation of *Z. tritici* was made from each leaf. But field collections made in California, Switzerland, Israel, and Uruguay often included isolations made from more than one discrete lesion on the same leaf or from more than one pycnidium in the same lesion. In these cases, more than one genotype per leaf was commonly found. For example, in the collections from Uruguay and Israel, where an average of 2 or 3 lesions were sampled per leaf, respectively, there was an average of 2 or 3 genotypes found per leaf, respectively. In one naturally infected Swiss field, microtransects were made through 5 lesions, with one lesion investigated per leaf, and each leaf coming from a different tiller growing in the same 1 m^2^ area. In this case, 15 different *Z. tritici* genotypes were identified among 158 isolates sampled from the 5 lesions, giving an average of 3 different genotypes per lesion (Linde et al. 2002). Our estimates of genotype diversity, focused on the most active phase of disease development and involving the four upper leaf layers, assumed that only one genotype is found on a leaf. We recognize that this is likely to underestimate the true number of genotypes because it is common to find more than one discrete STB lesion on a leaf and more than one genotype within a lesion, especially in fields experiencing moderate to high levels of infection.

**Table 1.** Number of unique *Zymoseptoria tritici* genotypes found on a determined number of leaves and lesions in 30 naturally infected wheat fields located on four different continents.

| Location (number of fields sampled) | Number of leaves | Number of lesions | Number of genotypes | Reference |
| --- | --- | --- | --- | --- |
| USA, California, Davis | 19 | 35 | 22 | McDonald and Martinez 1990 |
| USA, Oregon, Corvallis | 544 | 544 | 497 | Linde et al. 2002 |
| Switzerland (4) | 280 | 280 | 341 | Linde et al. 2002 |
| Israel, Nahal Oz | 47 | 135 | 158 | Linde et al. 2002 |
| USA, Texas (2) | 75 | 75 | 73 | Linde et al. 2002 |
| Argentina (3) | 193 | 193 | 154 | Juergens et al. 2006 |
| Uruguay, Colonia | 23 | 41 | 41 | Juergens et al. 2006 |
| Australia (4) | 193 | 193 | 176 | Juergens et al. 2006 |
| Tunisia (5) | 218 | 218 | 216 | Boukef et al. 2012 |
| Czech Republic (7) | 184 | 184 | 158 | Drabešová et al. 2013 |
| France, Grignon (2) | 680 | 680 | 656 | Morais et al. 2019 |
| Totals | 2456 | 2578 | 2492 |  |

To estimate the number of spores produced in a single *Z. tritici* pycnidium, we used published results coming from several labs over many years. All calculations were obtained from leaves infected under greenhouse conditions, in some cases from pycnidia located on the second or third leaves of seedling plants (e.g. Stewart et al. 2018) and in other cases from pycnidia located on the top leaves of adult plants (e.g. Suffert et al. 2013). The earliest work (Eyal 1971) marked individual pycnidia and carefully pipetted the entire mass of spores emerging as a cirrhus from each pycnidium over several wetting and drying periods to estimate the total number of spores coming from individual pycnidia over time. Most methods relied on counting the number of pycnidia in a group of leaf lesions (distinct or coalescing, depending on the STB severity) found on a whole leaf either manually (Gough 1978; Stewart et al. 2018) or by using automated image analysis (e.g. Yates et al. 2019), and then washing all spores off of that leaf after placing it into a humid environment that led to extrusion of cirri from all mature pycnidia. Another approach took into account the overall capacity of all pycnidia on a leaf to produce spores over a defined period of time, by washing spores from a leaf over several cycles of spore production (Suffert et al. 2013; Boixel 2020, p.229). Individual studies found differences in spore production among different *Z. tritici* strains (Gough 1978; Stewart et al. 2018; Suffert et al. 2013) and among different wheat cultivars (Eyal 1971; Gough 1978; Hess and Shaner 1987; Suffert et al. 2013). The total number of measurements made and the total numbers of pycnidia included in the measurements also differed widely among studies (Table 2). All told, data were obtained from 16 different cultivars that differed in their degree of STB resistance and included at least 45 strains of *Z. tritici* that differed in their overall reproductive capacity. The average values reported in different studies ranged from 830 to 11,500 spores per pycnidium. The mean value over all studies, weighted according to the number of measurements made (but excluding the values from Hess & Shaner 1987 because they did not report the number of measurements) was 5322 spores per pycnidium.

**Table 2.**
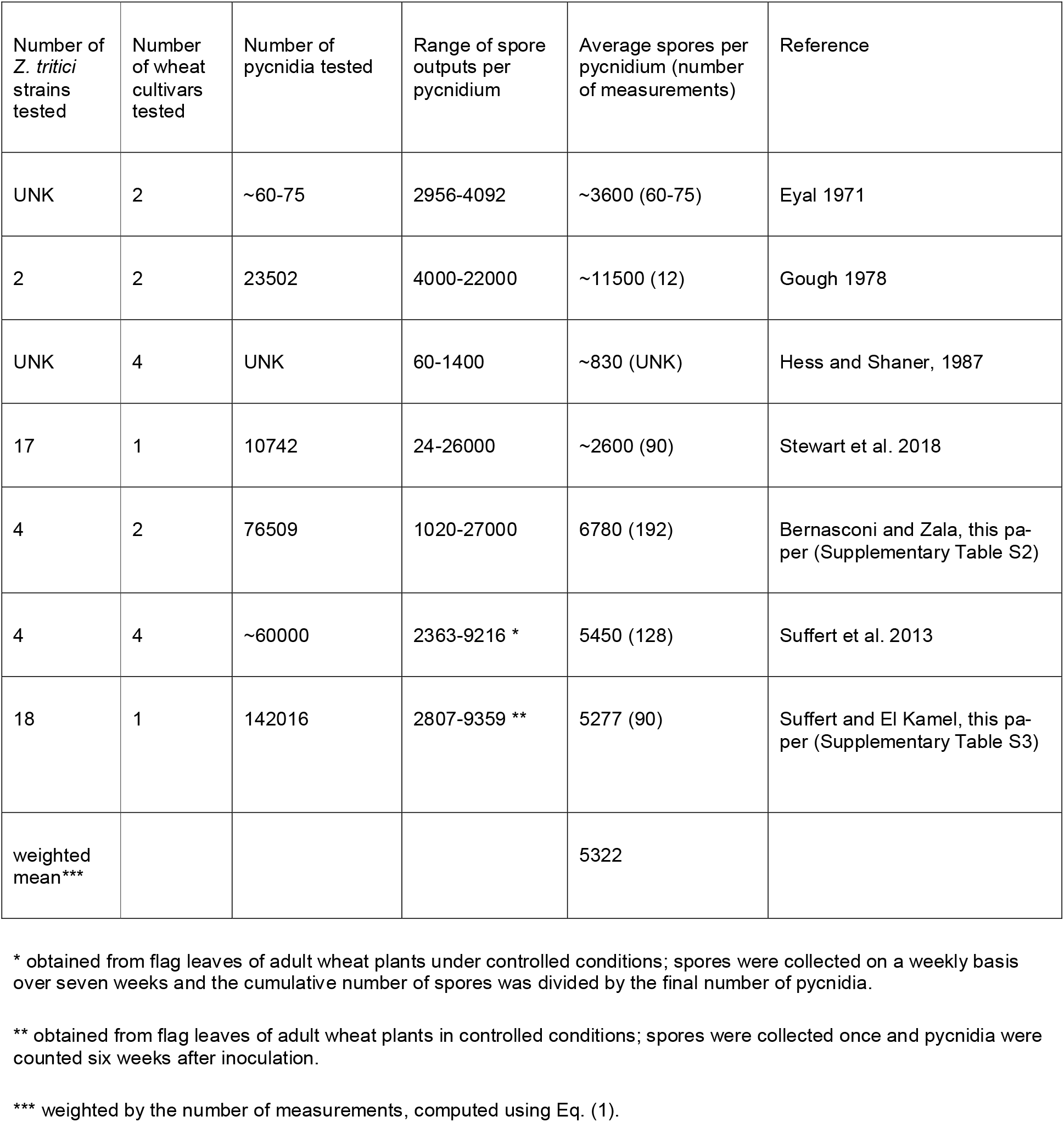
Average number of pycnidiospores produced within a *Zymoseptoria tritici* pycnidium. UNK = Unknown

| Number of *Z. tritici* strains tested | Number of wheat cultivars tested | Number of pycnidia tested | Range of spore outputs per pycnidium | Average spores per pycnidium (number of measurements) | Reference |
| --- | --- | --- | --- | --- | --- |
| UNK | 2 | ~60-75 | 2956-4092 | ~3600 (60-75) | Eyal 1971 |
| 2 | 2 | 23502 | 4000-22000 | ~11500 (12) | Gough 1978 |
| UNK | 4 | UNK | 60-1400 | ~830 (UNK) | Hess and Shaner, 1987 |
| 17 | 1 | 10742 | 24-26000 | ~2600 (90) | Stewart et al. 2018 |
| 4 | 2 | 76509 | 1020-27000 | 6780 (192) | Bernasconi and Zala, this paper (Supplementary Table S2) |
| 4 | 4 | ~60000 | 2363-9216 * | 5450 (128) | Suffert et al. 2013 |
| 18 | 1 | 142016 | 2807-9359 ** | 5277 (90) | Suffert and El Kamel, this paper (Supplementary Table S3) |
| weighted mean*** |  |  |  | 5322 |  |

\* obtained from flag leaves of adult wheat plants under controlled conditions; spores were collected on a weekly basis over seven weeks and the cumulative number of spores was divided by the final number of pycnidia.
\*\* obtained from flag leaves of adult wheat plants in controlled conditions; spores were collected once and pycnidia were counted six weeks after inoculation.
\*\*\* weighted by the number of measurements, computed using Eq. (1).

Estimates of the number of pycnidia found on a leaf came from only one lab that relied mainly on automated image analysis to identify and count pycnidia (e.g. Stewart and McDonald 2014; Stewart et al. 2016). These data were obtained from naturally infected leaves, with many coalescing lesions in cases of high STB severity, which made it impractical to estimate the number of pycnidia per lesion induced by a single genotype. Automated image analysis has been used in this lab for 10 years to identify and count *Z. tritici* pycnidia in the framework of experiments seeking to quantify differences in virulence and reproduction among strains of *Z. tritici* (Stewart et al. 2018; Dutta et al. 2021) and differences in quantitative STB resistance among wheat cultivars (Stewart et al. 2016; Karisto et al. 2018; Yates et al. 2019; Mikaberidze and McDonald 2020). These experiments were conducted mainly using seedling plants grown under controlled greenhouse conditions (e.g. Stewart et al. 2014), but several more recent experiments collected large datasets from naturally infected leaves of adult plants grown under field conditions (Stewart et al. 2016; Karisto et al. 2018; Mikaberidze and McDonald 2020). Estimates of pycnidia found within isolated lesions (a condition closer to what is observed in the field when STB severity is low) obtained by Morais et al. (2015) from flag leaves of adult plants inoculated with a low concentration of ascospores, showed that a lesion, whose size ranged from 10 to 20 mm^2^ (typically 1-2 mm wide x 10-20 mm long) contains approx. 20 to 40 pycnidia. However, we used here only the numbers of pycnidia found on naturally infected leaves taken from adult plants grown under natural field conditions. The numbers shown in Table 3 used automated image analysis to count a total of 2.95 million pycnidia on a total of 22,395 naturally infected leaves growing under field conditions over two years and in two locations (Stewart et al., 2016; Karisto et al., 2018). The weighted mean of 132 pycnidia per leaf was used for the calculations of spore production.

**Table 3.** Number of *Zymoseptoria tritici* pycnidia found per leaf in natural field infections using automated image analysis.

| Location, year | Number of wheat lines analyzed | Number of leaves analyzed | Range in pycnidia number per leaf | Mean number of pycnidia per leaf (± standard error) | Reference |
| --- | --- | --- | --- | --- | --- |
| Switzerland, 2015 | 39 | 733 | 40-326 | 148±11 | Stewart et al. 2016 |
| Oregon, 2015 | 9 | 240 | 52-1471 | 641±172 | Stewart et al. 2016 |
| Switzerland, 2016 t<sub>1</sub> | 334 | 10269 | 3-304 | 62±3 | Karisto et al. 2018 |
| Switzerland, 2016 t<sub>2</sub> | 335 | 11153 | 17-840 | 185±7 | Karisto et al. 2018 |
| Weighted mean * | | | | 132±4 | |

\* weighted by the number of leaves in each experiment, computed using Eq. (1), its standard error was computed using Eq. (2).

The leaf area exhibiting disease symptoms, measured at different spatial scales that can range from individual leaves or plants to entire crop canopies, is often used to estimate the intensity of a field epidemic, typically producing measures of disease incidence and/or severity. We estimated the percentage of diseased wheat leaves in natural STB epidemics characterized by low, moderate and high levels of disease intensity in order to cover a wide array of epidemic situations. We used datasets collected during 10 years (2008-2017) of field experiments conducted in Grignon, France. The experiments had three treatments corresponding to three different tillage and crop residue management practices representative of those practiced in the Paris basin, one of the main wheat producing areas in Europe (Suffert and Sache, 2011; Suffert et al. 2018). The datasets included measurements of disease incidence (the percentage of leaves carrying at least one STB lesion) and severity (the percentage of leaf area covered by STB lesions) for the three or four top leaf layers of wheat cv. Soissons (moderately susceptible to STB and among the most popular wheat cultivars in France from 1990 to 2015). These assessments were made several times during each growing season. The measurements were conducted visually on whole plants sampled in the field (25 plants per plot from 2008 to 2011 and 15 plants per plot from 2012 to 2017). For the analyses in the present study, we selected one date per year during the most active phase of disease development (centered around May: from late April to early June; see Supplemental Table 1). Selecting the final dates for assessment in each season (i.e., late June to early July) would lead to higher estimates of the final incidence and severity, but the pathogen population size at that stage would no longer be relevant for the further epidemic development during the same season. At earlier dates the epidemic development was more vulnerable to unfavorable weather conditions and measurements may not accurately represent the epidemiologically relevant pathogen population sizes. Mean values of incidence and severity were calculated with measurements averaged over the three different treatments and leaf layers. These values are shown in Figure 1. By conducting K-means clustering on these data, we found 18%, 44% and 83% STB incidence to represent the percentage of diseased leaves found in fields with low, moderate and high STB epidemic severity, respectively. These measures of disease incidence were used in the calculations of genotype diversity and spore production.

**Figure 1.**
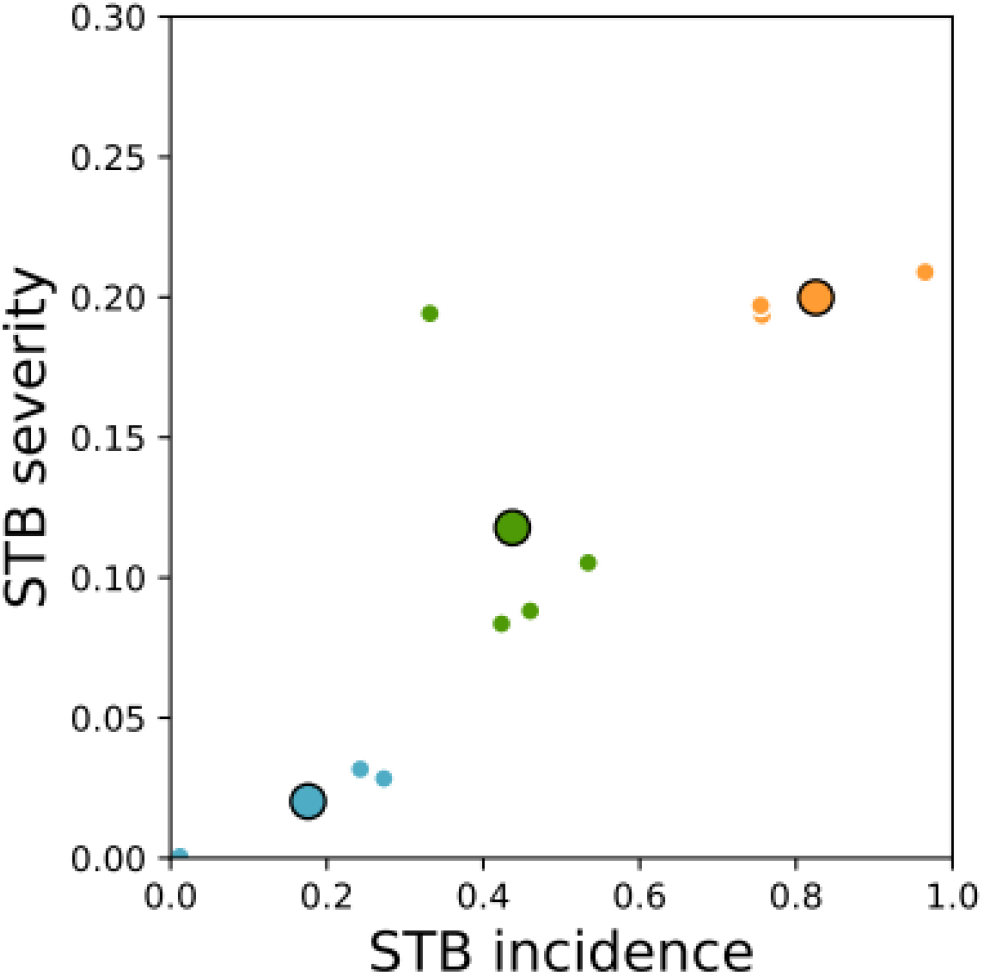
STB severity versus incidence measured in the field over 10 consecutive years (2008-2017; Suffert and Sache 2011; Suffert et al. 2018). Means over field assessments in each year are shown as small circles (for dates and values see Supplementary Table S1). Different colors correspond to three clusters obtained using K-means clustering: low epidemics (blue), moderate epidemics (green) and high epidemics (orange). Large circles show mean values within each cluster.

We computed global means over a number of different experiments to calculate the number of different genotypes per leaf (Table 1), the number of pycnidiospores per pycnidium (Table 2) and the number of pycnidia per leaf (Table 3) as weighted means:

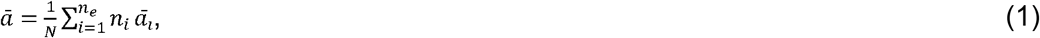

where *ā*_*i*_ is the mean value in experiment *i,n_i_* is the number of measurements conducted in experiment *i,n_e_* is the number of experiments, and *N* is the total number of measurements in all experiments. This way of computing global means incorporates the fact that different experiments had different numbers of measurements, and hence experiments with higher numbers of measurements should contribute more to the global mean values. Standard errors of weighted global means in Eq. (1) were computed as

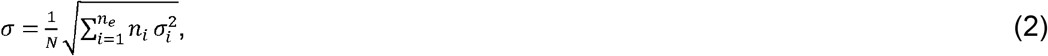

where *σ*_*i*_ is the standard error of the mean in the experiment *i*.

Based on our estimates of numbers of spores, we also estimated the numbers of adapted *Z. tritici* mutants produced during low, moderate and high epidemic years per hectare of wheat field. To obtain these estimates, we considered that the pathogen population undergoes *k* non-overlapping rounds of asexual reproduction (or generations) and estimated the number of spores *n*_*k*_ produced after *k* generations using the geometric growth model:

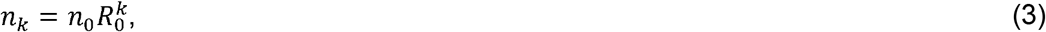

where *n*_*0*_ is the initial number of spores, *R*_0_ is the basic reproduction number of the pathogen, and *n*_*k*_ is the number of spores produced after *k* generations. The number of adapted mutants generated at *k*th round of reproduction is then

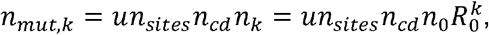

where *u* is the mutation rate per cell division per site, *n*_*sites*_ is the number of sites at which adaptive mutations can occur, and *n*_*cd*_ is the number of cell divisions per generation. To calculate the total number of adapted mutant spores *N*_*mut*_,_*k*_ produced in the course of *k* generations, we compute the sum of *n*_*mut,k*_ over *k*

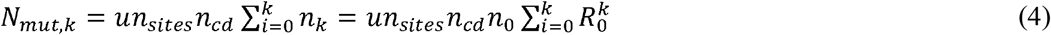

To estimate the number of adapted mutant spores we set the number of generations to *k*=3, use a conservative estimate of the basic reproduction number of *Z. tritici R*_0_=4 (Mikaberidze et al., 2017), a mutation rate per cell division per site of *u*=3 × 10^−10^ (Habig et al., 2021), and assume that there are 100 cell divisions per generation (*n*_*cd*_=100).

Data analyses were conducted using the Python programming language (v. 3.8.5). We quantified the correlation between the STB severity and incidence using Spearman’s correlation coefficient (“spearmanr” function in the scipy.stats package v. 1.5.2). We conducted K-means clustering of the severity/incidence dataset to characterize different degrees of STB epidemics (“Kmeans” function with three clusters in sklearn.cluster package v. 0.23.2).

## 3. Results

### 3.1. How many unique pathogen genotypes exist per hectare?

The STB assessments made during the most active phase of disease development from each of 10 years of field data from INRA Grignon allowed us to characterize the relationship between STB disease severity and the incidence of diseased leaves in natural epidemics. This relationship was close to linear and the two quantities exhibited a strong correlation (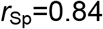, *p*=2.2 × 10^−3^; Figure 1). Based on these experimental findings, we assume the following typical values of STB severity and incidence corresponding to three classes of epidemics: low, moderate and high (with values corresponding to the large circles in Figure 1 showing means within the three clusters). In a low epidemic year (on average 2% of the leaf area covered by lesions; blue points), 18% of the leaves in an infected field had at least one STB lesion, while in a moderate epidemic year (12% of the leaf area covered by lesions; green points), 44% of the leaves were diseased, and in a high epidemic year (20% of the leaf area covered by lesions; orange points) 83% of the leaves were diseased.

Given the finding that each leaf is infected by a minimum average of one unique *Z. tritici* genotype in a naturally infected field (Table 1), and assuming that there are 18 million leaves per hectare, these rates of infection predict approx. 3.2, 7.9 and 14.9 million *Z. tritici* genotypes per hectare will be found in fields with low, moderate and high levels of infection, respectively. We consider it worthwhile to mention some caveats at this point. On the one hand, these numbers may over-estimate the true number of genotypes because the lesions present on the four upper leaves result from multiple cycles of asexual reproduction. The number of cycles of asexual reproduction occurring during an epidemic is estimated to range from 3-6 according to local climatic conditions (Lovell et al. 2004; Karisto et al., 2018). On the other hand, these numbers may under-estimate the true number of genotypes because there can be more than four leaves per tiller, an infected leaf often contains more than one discrete lesion and the majority of lesions contain more than one genotype (Table 1).

### 3.2. How many spores are produced per hectare?

The number of pycnidia forming on a leaf will be affected by the degree of host resistance, the reproductive capacity of the strain infecting each leaf, and the microenvironment in the leaf during the infection process. Using automated image analysis, it was possible to count large numbers of pycnidia on individual leaves of seedlings infected by particular pathogen strains under standardized greenhouse conditions. The average numbers of pycnidia per leaf following inoculation of 12 host genotypes by 145 global strains of *Z. tritici* (encompassing approx. 11,000 leaves) ranged from 81 to 377 on the three most susceptible cultivars and from 7 to 10 on the three most resistant cultivars tested under highly controlled greenhouse conditions (Dutta et al. 2021). It is more challenging to count pycnidia on naturally infected leaves growing under variable field conditions, but these data are available from field sites in Switzerland and Oregon, where measurements were made from 335 and 9 host genotypes, respectively (Table 3). In the Oregon field experiment, no fungicides were used and conducive environmental conditions enabled a high intensity of STB to develop, leading to an average of 641 pycnidia detected per leaf across 240 leaves. In the two years of the Swiss experiments, environmental conditions favored development of STB, but heavy fungicide applications suppressed the overall levels of STB, leading to a low-moderate level of infection overall, with an average disease rating at the final time point (approximately GS80) of only 2.5 on a 1-9 assessment scale. In these three leaf collections (22,155 leaves total), an average of 132 pycnidia were identified per leaf.

If we assume that only 18% of the upper four leaves are diseased in a low epidemic year, and that only 132 pycnidia form, on average, on these leaves, then we expect to find about 0.4 billion pycnidia per hectare. In a moderate epidemic year with 44% of leaves diseased, we expect to find approx. 1.0 billion pycnidia, and in a high epidemic year with 83% of leaves diseased we expect to find approx. 2.0 billion pycnidia per hectare. We anticipate that in some high epidemic years, more pycnidia will form on average on each leaf, as indicated by the 641 pycnidia found per leaf in the Oregon field trial. If we combine a high incidence of disease (83%) with a high number of pycnidia per leaf (641), then we could expect to find up to 9.6 billion pycnidia forming per hectare.

Under the assumption that an average of 5322 spores are produced per pycnidium, we estimate that approx. 2.3 × 10^12^ spores per hectare are produced in a low infection year (2% severity, 18% incidence), approx. 5.6 × 10^12^ spores per hectare are produced in a moderate infection year (12% severity, 44% incidence), and 10.5 × 10^12^ spores per hectare are produced in a high infection year (20% severity, 83% incidence). We note here that these should be considered average values of spore production that could be expected across a wide range of host cultivars, pathogen strains and environmental conditions.

If we combine the higher number of pycnidia found per leaf in Oregon (641) with the highest average number of spores found per pycnidium (11,500, Gough et al. 1978) and assume that 83% of leaves are diseased, we arrive at 10^14^ spores per hectare. If we use the highest reported values for each factor, with 97% of leaves infected (i.e., 17.5 × 10^6^ leaves per hectare), 1471 pycnidia per leaf, and 27,000 spores produced per pycnidium, we arrive at an upper estimate of approx. 7 × 10^14^ spores per hectare. Given these last two calculations, we consider it possible that 1-7 × 10^14^ spores per hectare may be produced on a susceptible cultivar grown without fungicide treatments in an STB-conducive environment (i.e. in a very wet year).

### 3.3. How many spores carry adaptive mutations per hectare?

We use our estimates of overall numbers of spores produced per hectare of wheat to estimate the number of *Z. tritici* spores that carry mutations potentially conferring adaptation to control measures such as fungicides or disease-resistant cultivars. We consider pathogen adaptation to azole fungicides targeting the sterol 14_α_-demethylase cytochrome P450 encoded by the *Cyp51* gene (Cools & Fraaije, 2013) and to STB resistance encoded by the *Stb6* resistance gene (Saintenac et al., 2018) in wheat as prime examples in our estimates. A number of mutations in the *Cyp51* gene of *Z. tritici* are known to confer varying degrees of resistance to azole fungicides (Cools et al., 2011). At least 130 different haplotypes of the *Cyp51* gene were identified encoding various combinations of 30 amino acid substitutions (Huf et al., 2018). Similarly, a number of mutations in *AvrStb6* allow the pathogen to avoid recognition by the *Stb6* gene product, thereby conferring an adaptive advantage (Brunner & McDonald, 2018; Stephenson et al., 2021). At least 44 different *AvrStb6* haplotypes were identified that encode various combinations of >60 amino acid substitutions (Stephenson et al. 2021). For both genes, however, different mutations are likely to confer different quantitative phenotypes due to the complex shape of the associated fitness landscapes (Cools et al., 2013). To take this into account, we assume that adaptive point mutations can occur at only five nucleotide positions in each of the two genes.

The number of adapted mutant spores was estimated using the geometric growth model [Eq. (3)]. According to Eq. (4), we estimated the total number of adapted mutant spores of *Z. tritici* produced during three generations that correspond to the decisive, final period of epidemic development as 2.8 × 10^7^ (low epidemic year), 7 × 10^7^ (moderate epidemic year) and 1.3 × 10^8^ (high epidemic year). These numbers should be considered conservative estimates, because *Z. tritici* is a multicellular fungus and the number of mitotic cell divisions occurring during its development *in planta* that contribute to mutations in spores can be much higher than 100, but it is difficult to estimate the total number of cell divisions occurring from the time a germinating spore successfully infects a leaf until a resulting pycnidium produces a new generation of spores. Hence the actual number of mutant spores may be considerably higher.

## 4. Discussion

According to our estimates, a typical wheat field infected by *Z. tritici* contains between 3-15 million unique *Z. tritici* genotypes per hectare, with an average of at least one unique pathogen genotype per diseased leaf. We estimate that this population will produce a minimum of between 2-11 trillion asexual spores per hectare. It is widely accepted that many plant pathogens produce large numbers of propagules during an epidemic and that many pathogen populations can carry a large amount of genetic diversity, but we are not aware of any other robust quantitative estimates for these parameters in the peer-reviewed literature. To obtain these estimates, we brought together a number of datasets collected over many years across a range of geographically diverse locations. These numbers highlight the remarkably high degree of genotype diversity and spore production associated with natural STB epidemics, and can be used to inform our understanding of pathogen adaptation to fungicides, disease resistance genes and changing environments.

### 4.1 The range of estimates indicates very high values for each parameter

We sought to obtain “average” values for each parameter used in our calculations, though we recognize that a wide range of values can be ascribed to each parameter due to differences in local conditions (e.g., in degrees of host susceptibility, pathogen virulence, and conduciveness of environmental conditions). We decided that hectares and annual cropping seasons are the most reasonable spatial and temporal units for measurements of genetic diversity and spore production in an agricultural pathogen because wheat is typically grown as annual monocultures in fields that cover a large surface area.

Wheat is grown globally, with different densities of leaves reflecting differences in local growing conditions. We chose an average value of 18 million leaves per hectare for all of our calculations, but our approach can readily accommodate higher or lower planting densities to prepare locally relevant estimates. Our assumptions of the proportion of diseased leaves in a field (incidence) came from 10 years of field data collected from only one cultivar growing in only one location. We recognize the added value of obtaining comparable estimates of incidence from a wider array of cultivars and growing environments and hope that these data will become available in the future. If they already are available for some locations, our approach can easily accommodate differences in disease incidence. Our estimates of the number of pycnidia formed on naturally infected leaves were obtained using a single technology (automated image analysis) and included leaves coming from only two locations and two years. Though we recognize the limitations inherent in these datasets, we think the estimates are reasonable given that they included counts of nearly 3 million pycnidia coming from a wide array of host and pathogen genotypes. The average value used in our calculations also fell within the range of estimates coming from greenhouse experiments conducted under highly controlled conditions using 12 host lines and 145 global pathogen genotypes (Dutta et al. 2021), increasing our confidence in using this value. Our estimates of the number of spores produced per pycnidium were based on several different approaches and were obtained from several labs over many years. Though the range of reported values was quite wide, we feel reasonably confident that our weighted average value of 5322 reflects the typical number of pycnidiospores per pycnidium likely to be found across a wide variety of cultivars, pathogen strains and field environments.

We brought together many studies of genotype diversity based on >2400 genotyped *Z. tritici* individuals collected in naturally infected wheat fields from around the world (Table 1). Based on these analyses, we estimated that a typical field infected by *Z. tritici* carries between 3.2-14.9 million unique *Z. tritici* strains per hectare. We consider these to be conservative estimates. Actual numbers could be 2-3 times higher given that most studies of genotype diversity included only one strain per leaf, while studies that included more than one strain per leaf typically reported more than one unique genotype per diseased leaf (Linde et al., 2002; Juergens et al. 2006).

Bringing together the average values of leaf density, STB incidence, pycnidia per leaf and spores per pycnidium allowed us to estimate an average number of *Z. tritici* pycnidiospores likely to be produced during an epidemic under low, moderate and high disease intensities. Our calculations suggest that a typical field infected by STB will produce between 2.3 and 10.5 trillion pycnidiospores per hectare, though a heavily infected field may produce as many as 700 trillion spores per hectare. It is important to note here that only a small fraction of these spores will actually contribute to each infection cycle because a great majority of the generated propagules fail to cause new infections for a variety of reasons, e.g. the spores do not land on host tissues, are degraded or non-viable, fail to penetrate the epidermis, etc. But these spores contain a vast reservoir of mutations on which selection may operate as described below.

### 4.2 What is the source of the high genotype diversity found in *Z. tritici* populations?

To answer this question we can draw on recent, detailed analyses of whole genome sequences that were conducted in *Z. tritici* (Hartmann et al., 2017; Singh et al., 2021; Plissonneau et al., 2018). Among 107 strains from a global collection, about 0.78 million high-confidence single-nucleotide polymorphisms (SNPs) were detected with an average of 20 SNPs per kilobase (Hartmann et al., 2017). Among 177 isolates collected from the same wheat field in Switzerland, about 1.5 million high-confidence SNPs were detected with an average of 37 SNPs per kilobase (Singh et al., 2021). While SNPs seem to dominate the genetic variation in *Z. tritici* populations, structural polymorphisms resulting from chromosomal rearrangements (Goodwin et al., 2011; Croll et al., 2013) and gene gain/loss (Hartmann and Croll, 2017; Plissonneau, 2018; Badet et al. 2020) also make substantial contributions to genotype diversity. Considered together, these studies suggest that a substantial portion of the genetic diversity observed in *Z. tritici* populations results from random mutations that produce polymorphisms within local populations. These mutations can persist within local populations as a result of the large population sizes associated with the estimates of spore production presented here and the high effective population sizes estimated in earlier studies (Zhan et al. 2001; Zhan and McDonald 2004; Singh et al., 2021). Sexual recombination does not influence nucleotide diversity, but it does increase genotype diversity via formation of new combinations of alleles of different genes and through intragenic recombination. *Z. tritici* has a mixed reproductive system (Chen and McDonald 1996; Zhan et al., 1998): within-season epidemics are driven by multiple rounds of asexual reproduction via pycnidiospores, while the initial inoculum in the following season is dominated by sexual ascospores (Shaw and Royle, 1989; Suffert and Sache, 2011; Suffert et al., 2011; 2018). Despite a likely bottleneck in pathogen population size in the absence of living wheat plants between wheat growing seasons, the transition from one season to the next is expected to re-establish high levels of genotype diversity because ascospores resulting from sexual recombination initiate the STB epidemic each season. Additional factors that contribute to high genotype diversity are new cycles of ascospore production within infected fields during the growing season (Zhan et al. 1998), high gene flow among fields (Boeger et al. 1993) enabled by long-distance dispersal of the sexual ascospores (Shaw and Royle, 1989), and the pathogen’s ability to persist on infected wheat debris between growing seasons (Suffert and Sache, 2011; McDonald and Mundt 2016). The recent discovery that *Z. tritici* forms chlamydospores suggests an additional contribution from a long-lived spore bank in the soil that may serve as a long-term reservoir of genetic diversity (Sardinha-Francisco et al., 2019). Thus, the high genotype diversity found in *Z. tritici* populations is consistent with the current knowledge of the population biology and population genomics of this pathogen.

### 4.3 How do *Z. tritici* populations maintain high genotype diversity under strong selection?

In wild plant pathosystems, local genetic and environmental heterogeneity of host plant populations contributes to maintenance of genetic diversity in the corresponding pathogen populations (Laine et al., 2011). In contrast, agricultural ecosystems such as those planted to wheat are typically grown in large, environmentally uniform fields consisting of genetically identical hosts. These wheat fields are often planted to disease-resistant cultivars and sprayed with fungicides to control diseases. Hence, we might expect that the uniform environment coupled with strong selective pressures will rapidly diminish the degree of genetic diversity found in *Z. tritici* populations. How do *Z. tritici* populations maintain such a high degree of genotype diversity over time and space under such highly selective conditions? Fungicides and disease-resistant wheat cultivars are not always expected to impose directional selection that will diminish genetic diversity in pathogen populations. In cases when a large and constant selective advantage is conferred by a single mutation, selection is indeed directional (e.g., resistance to quinone outside inhibitor (QoI) fungicides; Torriani et al., 2009; Estep et al., 2015). In contrast, when pathogen adaptation occurs via accumulation of several mutations, with each having a moderate and/or additive effect, selection can generate more complex and variable fitness landscapes associated with individual mutations and their combinations. Examples include adaptation of *Z. tritici* to azole fungicides (Estep et al., 2015) and to the disease resistance gene *Stb6* in wheat (Brunner et al., 2018): both the azole target site gene *Cyp51* and the avirulence gene *AvrStb6* exhibit substantial polymorphisms in *Z. tritici*, with population genetic analyses providing strong evidence of intragenic recombination and diversifying selection (Estep et al., 2015; Brunner et al., 2018). An analysis that included only recently collected *Z. tritici* populations found evidence for directional selection favoring a particular *AvrStb6* allele with a global distribution that likely emerged through convergent evolution in different populations. The same study showed that high levels of nucleotide and amino acid diversity were maintained for this gene in most sampled populations (Stephens et al., 2021). Additional selective factors that may maintain genotype diversity in *Z. tritici* include tradeoffs among life history traits (Dutta et al., 2021) and differential fitness of strains when they co-infect and co-exist within the same leaf (Linde et al., 2002; Zala et al., 2021).

### 4.4 Overall implications for pathogen adaptation

While the total number of spores produced during an epidemic has significant consequences for epidemic development, more important for long term evolutionary processes is the effective size of a pathogen population and the ability of this population to generate genotypes that are especially well adapted to the local environment, which may include resistant hosts and fungicides. Our estimates of the number of different genotypes include those resulting from sexual reproduction occurring between and within growing seasons within regions and those resulting from mutations that occur within fields. As the number of genotypes in a field increases, the probability that a genotype occurs that is able to overcome an STB control strategy implemented in that field (e.g. deployment of fungicides or resistant cultivars) also increases. Paradoxically, as the total number of genotypes increases, it becomes more difficult for a particular genotype to become dominant. The consequences of the pathogen population diversity for local adaptation also depend on the ability of adapted genotypes to persist over time (i.e. across asexual and/or sexual generations) and to be transmitted over larger spatial scales (i.e. across an entire field and/or between fields). A well-adapted genotype must first multiply locally and then be transmitted over a long distance in order to make a significant contribution to an epidemic. Then it must persist between growing seasons to contribute to future epidemics. These processes are considered in the remainder of this section.

The high genetic diversity found in wheat fields infected by *Z. tritici* is expected to drive a field population’s adaptation to local control measures and environmental conditions. Using the example of population adaptation to azole fungicides and the *Stb6* resistance gene in wheat, if we assume three generations of pathogen reproduction during the final phase of a cropping season, we estimate that there will be 28 million, 70 million and 130 million adapted mutant spores appearing in every hectare experiencing low, moderate and high epidemic intensity, respectively. Can these adapted mutants be maintained in a field population as standing genetic variation even in the absence of a specific selection pressure (i.e., deployment of an azole fungicide or a wheat cultivar with the *Stb6* resistance gene), or are they more likely to emerge *de novo* following the deployment of the selection pressure? To address this question, we would need to estimate the proportion of mutant spores that land on wheat leaves and are able to cause sporulating lesions. These estimates can then be incorporated into a population dynamical mathematical model that takes into account demographic stochasticity and the associated probability of stochastic extinction. However, even before such models are developed, these numbers (between 28 and 130 million adapted mutants per hectare per season) appear to be high enough to allow the adapted mutants to escape stochastic extinction and to be maintained in any field population even in the absence of the specific control measure. The mechanism of such maintenance can be mutation-drift equilibrium if the mutants are neutral or mutation-selection equilibrium in case the mutations are slightly deleterious in the absence of the control measure. These considerations lead us to conclude that resistance to fungicides and virulence to major resistance genes are likely to emerge from standing genetic variation in the majority of wheat fields.

We expect that any mutants conferring fungicide resistance or virulence against a given resistance gene will first be selected during the growing season and increase in frequency in treated fields via mostly asexual reproduction. At the next stage the selected strains will undergo mating and sexual recombination with a diverse pool of wild-type or mutated pathogen individuals and thereby embed the selected mutant alleles into diverse genetic backgrounds. These regular cycles of recombination make the adapted pathogen subpopulation carrying the mutation more robust with respect to changing environmental conditions and enables the long-term persistence of the adapted mutation. In a final step, any fitness penalties associated with the mutant alleles are likely to be ameliorated through compensatory mutations occurring either in the same gene or in unlinked genes that can be brought together by recombination.

### 4.5. Implications for field-scale strategies of disease management

Current epidemiological/evolutionary models that inform management of fungal diseases of crops, including STB, typically consider only two pathogen strains, a wild-type and an adapted strain (e.g. a fungicide-resistant or a virulent strain) that reproduce asexually (e.g. van den Bosch & Gilligan 2008; Lo Iacono et al., 2013; Mikaberidze et al. 2014). One conclusion from these models that is relevant to STB management is that in cultivar mixtures consisting of disease-resistant and disease-susceptible cultivars, higher proportions of a resistant cultivar make the mixture more durable with respect to breakdown of host resistance genes (Lo Iacono et al. 2013). Furthermore, studies that specifically parameterized these modeling approaches to *Z. tritici* populations concluded that lower fungicide doses slow down the emergence of fungicide resistance (Hobbelen et al., 2014; Mikaberidze et al., 2017), and that fungicide mixtures are typically more durable with respect to pathogen evolution compared to fungicide alternations when a high-risk fungicide is mixed with a low-risk fungicide (Elderfield et al., 2018). However, as we argue above, the high standing genetic diversity in *Z. tritici* field populations combined with large local populations and abundant sexual recombination represent key unconsidered factors that are likely to affect the emergence and persistence of adapted pathogen strains. If adapted mutants indeed pre-exist in local populations as part of standing genetic variation, this would alleviate the detrimental role of stochastic extinctions in the emergence of adapted pathogen strains (Lo Iacono et al. 2013; Mikaberidze et al. 2017). Moreover, as we explained above, a quick reshuffling of alleles via sexual recombination would embed the adapted mutants into diverse genetic backgrounds, making them more robust with respect to environmental stochasticity, which may constitute a key factor in their long-term persistence. These insights call for a re-consideration of mathematical modeling approaches that currently inform management of STB, because incorporation of the above factors may put the current conclusions into question.

Our findings also raise questions regarding the strategy of planting mixtures of wheat cultivars carrying different *Stb* resistance genes (Kristoffersen et al. 2020) to manage STB. We suggest that increasing the diversity of host populations at the field scale can have both beneficial and deleterious impacts in the presence of such highly diverse pathogen populations. For example, consider a field planted to a mixture of a susceptible wheat cultivar and a resistant cultivar carrying a fully effective *Stb* resistance gene with properties like *Stb6*. A mutation occurring in the larger pathogen population existing on the susceptible cultivar could confer virulence on the *Stb* gene found in the resistant cultivar. In a homogeneous population of susceptible cultivars, this mutant virulence allele exists as standing genetic variation, but would be unlikely to increase in frequency because it would not provide a competitive advantage. But in a mixture with the resistant cultivar, spores carrying this mutant allele are more likely to land on the nearby resistant cultivar where they can cause infection and multiply in the absence of competition from the more common avirulent strains found on the susceptible cultivar. In this way, cultivar mixtures can drive the emergence of new virulence alleles and facilitate the process of breaking resistance genes. In contrast, in a resistant pure stand, the number of new virulent mutant strains is expected to be lower because the total number of spores produced will be lower, but when virulent mutants emerge or are introduced into the field through gene flow, they will undergo stronger selection than in a mixture and will likely increase in frequency more rapidly in the pure stand.

However, cultivar mixtures can still effectively control STB and limit the loss in effectiveness of a resistance gene — in opposition to what we just described — when these mixtures are composed of cultivars carrying already broken *Stb* resistance genes (i.e. *Stb* genes whose corresponding virulence mutations are already present but do not yet occur at a high frequency in the pathogen population). A recent field experiment suggested that a mixture can maintain the efficacy of the resistance encoded by *Stb16q* through a decrease in the frequency of virulent strains infecting the susceptible cultivar and an increase in the frequency of avirulent strains occurring on the cultivar carrying *Stb16q* in the mixtures compared to pure stands. The observed changes resulted (i) on the one hand from virulence selection/counter-selection driven by exchanges of splash-dispersed asexual spores between cultivars depending on their respective proportions in the mixture (Orellana-Torrejon et al., 2022a), and (ii) on the other hand from sexual reproduction between virulent strains and avirulent strains that land on the cultivar carrying *Stb16q* and then recombine with virulent strains without the need to infect host tissues (Orellana-Torrejon et al., 2022b). This mechanism that explains the persistence (or even a slight increase) of avirulent strains in mixtures was experimentally established by Orellana-Torrejon et al. (2022c), who showed that symptomatic asexual infection is not required for a strain to engage in sexual reproduction [a similar finding was also reported for the *Stb6-AvrStb6* interaction (Kema et al., 2018)]. While cultivar mixtures can thus be an effective strategy to extend the useful life of already-defeated resistance genes or to enable recycling of defeated resistance genes, we should be aware, as previously stated, that cultivar mixtures including fully susceptible cultivars may also facilitate the emergence of virulent mutants that can overcome newly-deployed resistance genes. Following this line of reasoning, the benefit of mixtures vs pure stands would depend on the frequency of virulent strains in a local population, which in turn reflects the commercial life cycle of the corresponding varieties. The optimum strategy to formulate cultivar mixtures may be to mix varieties carrying quantitative resistance with varieties containing broken qualitative resistance (e.g. *Stb6*), while avoiding the use of fully susceptible varieties (to improve efficacy) and the introduction of new varieties containing unbroken qualitative resistance (to improve durability).

## Acknowledgements

Marcello Zala and Ons El Kamel assisted AB and FS, respectively, with generating the two previously unpublished spore count datasets shown in Supplementary Tables S2 and S3. AM thanks Daniel Croll for fruitful discussions.

---

**Supplementary Table S1.** Means over field assessments of STB incidence and STB severity measured at ten dates over ten consecutive years presented in Figure 1 of the main text (Suffert and Sache 2011; Suffert et al. 2018).

| Date | STB incidence | STB severity |
| --- | --- | --- |
| 2008-05-21 | 0.46 | 0.09 |
| 2009-05-27 | 0.42 | 0.08 |
| 2010-06-03 | 0.01 | 0.00 |
| 2011-04-28 | 0.27 | 0.03 |
| 2012-05-30 | 0.76 | 0.19 |
| 2013-05-22 | 0.33 | 0.19 |
| 2014-05-27 | 0.75 | 0.20 |
| 2015-06-04 | 0.97 | 0.21 |
| 2016-05-26 | 0.53 | 0.11 |
| 2017-05-22 | 0.24 | 0.03 |

## Notes

### Competing Interest Statement

The authors have declared no competing interest.

